# Engineered MS2 Virus Capsids for Cellular Display of Peptide Antigens

**DOI:** 10.1101/2025.08.19.671134

**Authors:** Hannah S. Martin, Paul Huang, Ian C. Leifer, Preeta Pratakshya, Matthew B. Francis

## Abstract

Our ability to respond to emerging pandemics and pathogen resistance relies critically on our ability to build vaccines quickly and efficiently. In this report we used an efficient enzymatic oxidative coupling reaction to create a viral capsid-based vaccine platform that is modular and quickly adaptable for many different pathogens. Tyrosinase-mediated oxidative coupling was used to conjugate C-terminal tyrosine residues on peptide antigens to cysteine residues installed inside MS2 viral capsids. This strategy is particularly promising because the capsids protect the internally conjugated peptides from protease degradation before they are delivered into cells. The vaccine constructs were tested for MHC presentation followed by T-cell activation. Mutants of the MS2 capsid itself activated DC2.4 cells, serving as an adjuvant to help induce the immune response to delivered antigens. The MS2-peptide constructs were shown to be stable in serum, activate DC2.4 cells, and lead to MHC-presentation of peptide antigens with subsequent activation of antigen-specific T-cell hybridomas. Taken together, these results demonstrate effective activation of the adaptive immune system *in vitro*. This synthetic platform can be used to build new vaccines for many different diseases for which immunodominant peptide antigens are known because the antigens can be quickly interchanged while the MS2 scaffold remains the same. Additionally, this platform allows for multiple peptide antigens to be delivered simultaneously in each capsid, which could provide enhanced immunity against resistant strains and be useful for cancer vaccine development.

## INTRODUCTION

Increases in global travel have rendered the spread of disease faster than ever, spurring the need to generate new vaccines at a rapid pace.^1^ While the traditional vaccine approaches of pathogen attenuation and inactivation are effective for producing strong immune responses, these approaches take many years to develop for each new pathogen.^2^ As a newer technology, subunit immunization approaches that only deliver small pieces of pathogens are emerging as safe and effective methods for creating vaccines very quickly.^2,3^ The antigenic segment used in subunit vaccines is carefully chosen from studies that show which components are primarily responsible for the immune response against a given pathogen.^4^ Peptides are one type of antigen of particular interest, as they represent only the amino acid sequence that is presented on major histocompatibility complex (MHC) molecules. However, small peptides typically do not lead to successful immunizations on their own because they are readily degraded in the blood and show poor uptake into antigen presenting cells. As a result, additional immune activation (usually an adjuvant) is needed to stimulate an immune response to them.^5,6^ This situation can be improved using delivery vehicles that exhibit increased cell uptake and (ideally) protect peptide antigens from degradation before they can be displayed.

Synthetic protein particles based on viral capsids, also known as virus-like particles (VLPs), are established platforms for delivering peptides to antigen-presenting cells. These constructs are typically made by fusing a desired sequence to the N or C terminus of the individual capsid monomers, but it can be challenging to find a location for peptide insertion that does not interfere with capsid assembly or alter its physical and biodistribution properties. As an alternative approach, one could consider the attachment of antigenic peptides to preassembled capsid particles using bioconjugation chemistry. Researchers have indeed conjugated antigens to VLPs using SpyCatcher/SpyTag systems and copper-catalyzed click chemistry as successful vaccination strategies.^7–10^

Herein we report an efficient chemical strategy to convert the protein capsid of bacteriophage MS2 into a peptide vaccine platform that could be used for many different antigen sequences. This protein nanocage consists of 180 identical coat protein monomers that self-assemble into *T* = 3 icosahedral structures with 32 pores, each 2 nm in diameter.^11,12^ These 2 nm pores are ideal for loading small molecule and peptide cargo because such species can passively diffuse through and be covalently attached to the *insides* of the capsids without requiring a disassembly/reassembly protocol.^12^This approach was demonstrated for the installation of up to 180 peptides on the inside surfaces of each capsid, thus minimizing the changes in the overall charges and solubilities of the particles that would be expected for external modification strategies. Moreover, proteases that could prematurely degrade the peptides would be expected to be too large to fit through the 2 nm pores. In previous studies we have shown that MS2 has prolonged *in vivo* circulation, with 10-15% of the injected dose per gram remaining in the blood after 24 h, observed in mice.^13^ While the wild-type MS2 capsid is not readily taken up into mammalian cells, our lab has recently reported engineered MS2 mutants that are endocytosed with much greater efficiency, potentially offering improvements in antigen display.^14^ Taken together, these features suggest that engineered MS2 viral capsids could provide a promising delivery system for peptide subunit vaccines.

## RESULTS AND DISCUSSION

### MS2 PKR Mutants show enhanced uptake by DC2.4 cells and promote dendritic cell activation

The primary role of a delivery vehicle is to take cargo into the cell, but wild-type MS2 (MS2 wt) capsids are normally taken up into mammalian cells at very low levels. To address this, our lab has previously developed mutants of MS2 that exhibit greatly increased endocytotic uptake.^14^ Two positively charged mutations, T71K and G73R (MS2 KR), were incorporated around the capsid pores (Figure 1a). These mutations create local positive charge regions around the capsid pores, while maintaining the net negative charge on the full particles. This lowers the risk of toxicity that is often seen in highly cationic delivery vehicles.^15–17^ The KR mutations are believed to bind to heparan sulfate molecules on the surface of mammalian cells, which aids in their uptake. Additionally, an N87C (C87) mutation provides an internal nucleophilic cysteine for bioconjugation of small molecule or peptide cargo that MS2 capsids with KR mutations can carry into cells.

**Figure 1.**
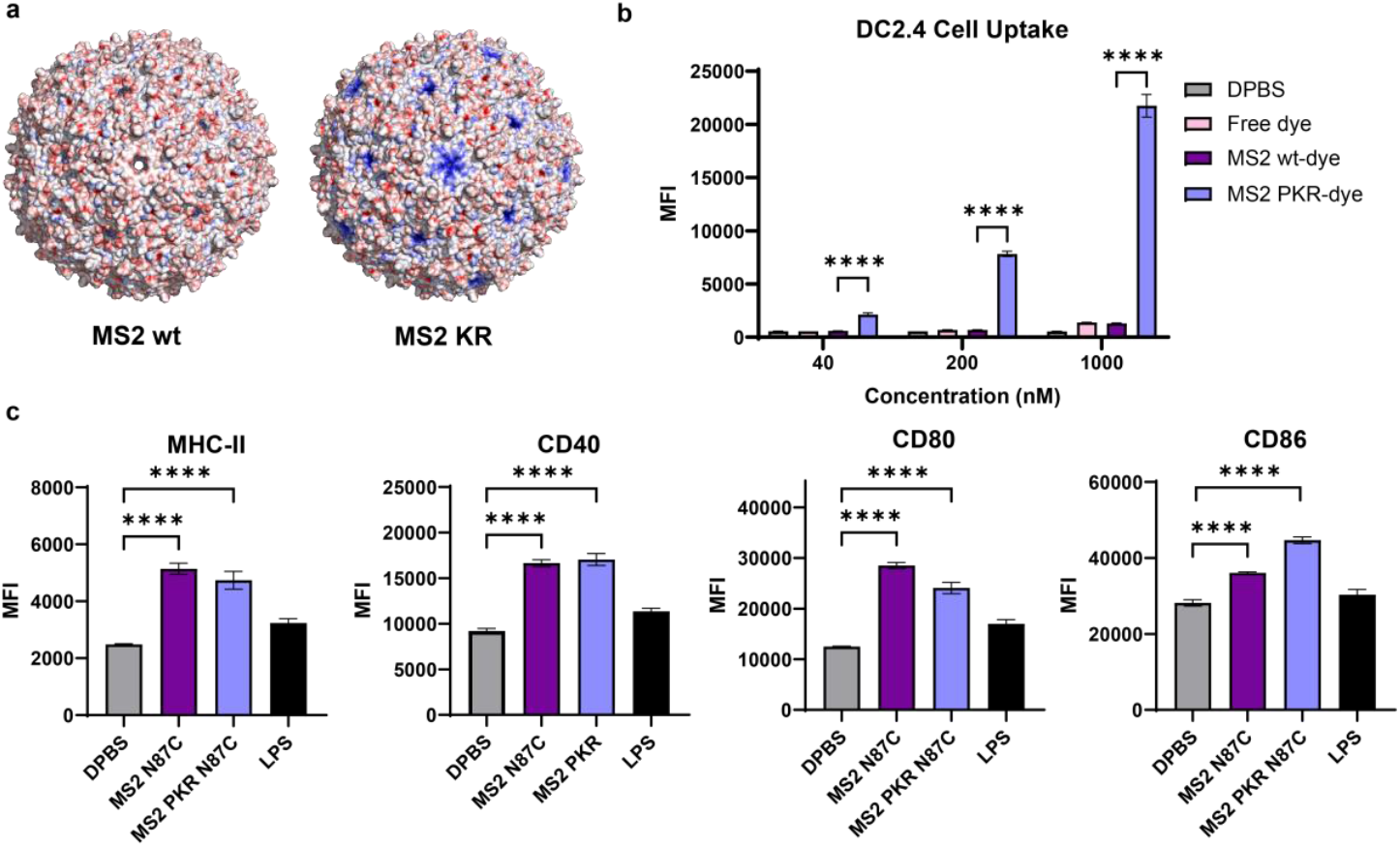
MS2 cell uptake and DC activation, a) Electrostatic surface potential maps of the wild type (MS2 wt) bacteriophage MS2 capsid and MS2 with T71 K. G73R (MS2 KR) mutations (both shown in blue). Positive charges are shown in blue and negative charges are shown in red. Capsids were rendered in PyMol v3.1.6.1 using the APBS (Adaptive Poisson-Boltzmann Solver) plugin, b) Cell uptake of MS2 capsid mutants in DC2.4 cells, c) Flow cytometry data of dendritic cell activation after treatment of DC2.4 cells with 1 pM of MS2 or 1 ng of LPS. N=3 biological replicates are shown.

Another MS2 mutation used in these studies is S37P, which has been shown to decrease the size of wild-type MS2 capsids from 27 nm to 17 nm in diameter.^18^ However, when the S37P mutation is combined with the KR mutations (MS2 PKR), the capsids maintain a 27 nm average diameter like the wildtype capsid (SI Figure S2).^19^ After internal labeling with AlexaFluor-594 maleimide on C87, MS2 PKR capsids showed substantially increased cell uptake into DC2.4 dendritic cells compared to wild type MS2 N87C (Figure 1b). The same trend was observed for U251MG glioma cells (SI Figure S3).

In addition to their enhanced cell uptake ability, MS2 capsids activated dendritic cells. To measure dendritic cell activation, DC2.4 cells were used. DC2.4 cells are an immortalized dendritic cell line that originated from C57BL/6 mouse bone marrow. These cells are antigen-presenting cells and express normal dendritic cell markers including CD11c and MHC-I molecules. DC2.4 cells naturally exist in an immature DC phenotype with low expression of MHC-II molecules and CD40, CD80, and CD86 costimulatory molecules.^20^ DC2.4 cells are also known to mature with certain stimuli, leading to the upregulation of MHC-II, CD40, CD80, and CD86. ^21–23^ To determine if MS2 can activate DCs, DC2.4 cells were treated with each MS2 mutant for 24 h followed by flow cytometry to measure the expression levels of four DC activation markers (MHC-II, CD40, CD80 and CD 86) after staining with the appropriate antibodies (Figure 1c). A low dose of 1 ng of LPS per well was used for the LPS controls.

Both MS2 wt and MS2 PKR capsids greatly elevated the expression of all markers compared to the DPBS control. Despite the differences in cellular uptake, both MS2 species resulted in higher DC activation across all four markers. The DC activation observed is likely partly a result of the endotoxin (LPS) present on the capsids from their recombinant expression in *E. coli*. Additionally, MS2 is known to encapsulate its own genetic material as well as other nucleic acids present in the expression system.^24^ These nucleic acids could be activating tolllike receptor pathways, which is commonly observed with immunization using VLPs.^25,26^ For use in vaccines, the DC activation caused by the capsids could be advantageous, serving as an adjuvant to stimulate an immune response during vaccination.

### Efficient conjugation of peptide antigens to MS2 using tyrosinase oxidative coupling

To modify the MS2 capsids, engineered interior C87 sites were used as nucleophiles. This allowed for up to 180 copies of a given peptide to be loaded inside each carrier. While maleimides are commonly used to couple cargo to cysteine-containing proteins, maleimide addition products can be reversible in serum,^27,28^ and such strategies would require the installation of a non-native functional group on the peptides. Using a tyrosinase enzyme from *Bacillus megaterium* (megaTyr), a covalent linkage can be formed between tyrosine and cysteine residues, two natural amino acids, via an oxidative coupling reaction.^29,30^ This linkage adds very minimal bulk and is irreversible in serum. Additionally, megaTyr can only oxidize terminal and highly surface-exposed tyrosine residues because the active site of the megaTyr enzyme is sterically occluded; thus, it does not react or interfere with internal protein tyrosines.^29^ The peptide antigen only needs to possess a tyrosine residue at the terminus to become a substrate for this reaction. Using tyrosinase to modify MS2 with peptides, different antigens can be quickly interchanged to create vaccines for a wide range of disease targets.

Tyrosine-containing peptide antigens were combined with MS2 PKR and reacted with megaTyr, which quickly formed the desired MS2-peptide conjugates with high labeling efficiency (Figure 2a, b). Three commonly studied model peptides for MHC presentation, Ova_257-264_ (Ova-I), Eα_52-68_ (Eα), and Ova_323-339_ (Ova-II) all coupled quite readily. Additionally, three peptides that are therapeutically relevant for targeting cancer, EphA2_883-891_ (EphA2), hTERT_613-626_ (hTERT), and Trp1_113-126_ (Trp1), were successfully coupled as well (Figure 2d).^31–33^ It is notable that the Trp1 peptide contains an internal disulfide that was not disrupted by the tyrosinase conjugation chemistry, as shown by mass spectrometry. This strategy can also be used to conjugate two different peptide antigens to the same capsid with controllable conjugation efficiency (SI Figure S4). Each peptide reacted with varying efficiency; most notably, the two peptides with higher pI values, hTERT and TRP1-disulfide, reacted less efficiently, but the amount of peptide used in a conjugation reaction can be increased to achieve higher % conjugation if desired.

**Figure 2.**
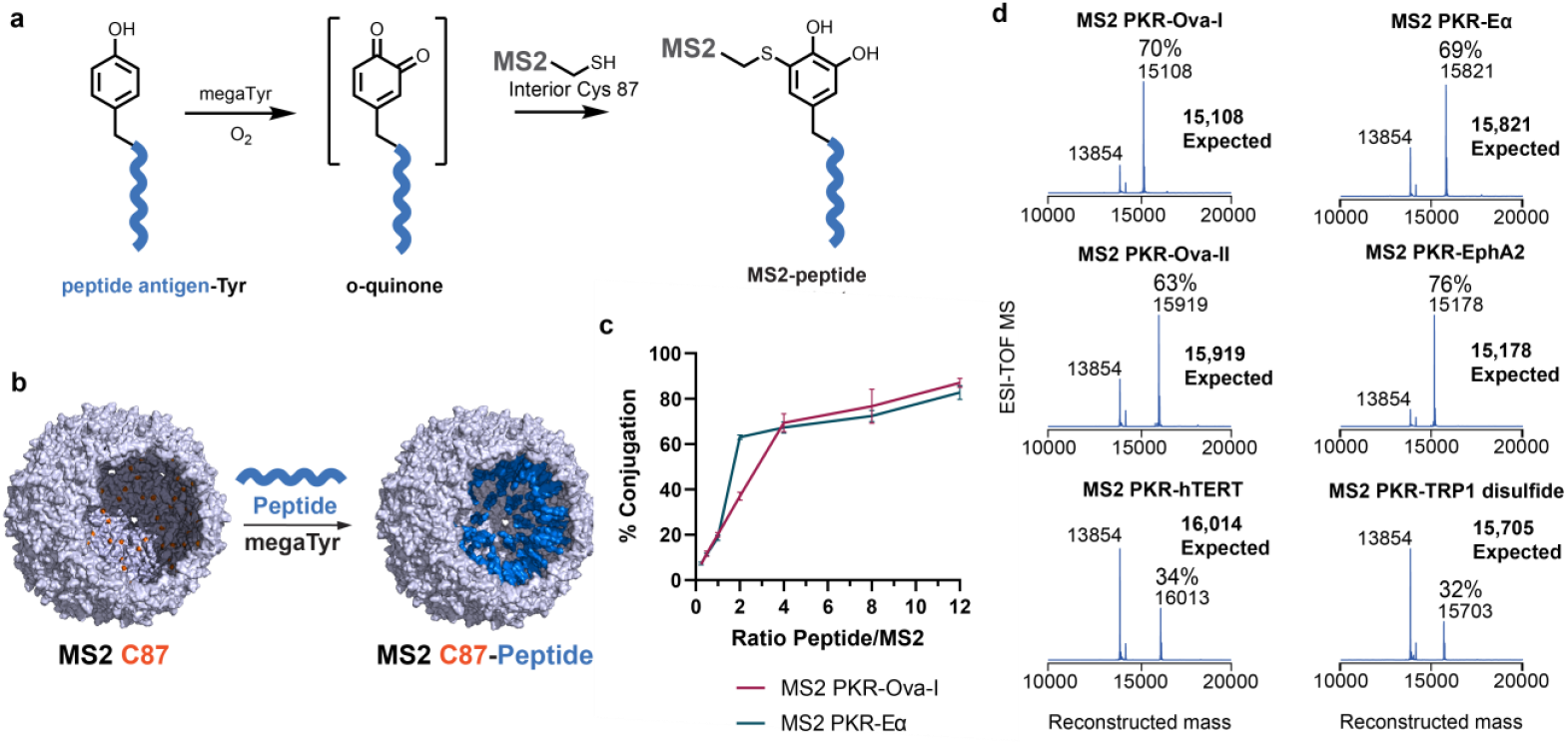
Peptide antigen conjugation to MS2. a) Tyrosinase-mediated oxidative coupling reaction of a tyrosine-tagged peptide antigen to the internal C87 mutation on MS2 PK.R. b) Structure of MS2 with a cut-out showing internal C87 (in orange) followed by internal coupling of Ova-I peptides (in blue), c) Peptide coupling yields for Ova-I and Ea attachment to MS2, as estimated using ESI-TOF MS. N=3 biological replicates are shown, d) Variety of MHC-1 and MHC-II peptide antigens conjugated to the internal C87 mutation of MS2 PK.R. The MS2 PK.R-TRP1 expected mass assumes the intact TRP1 disulfide.

MS2-peptide conjugates were primarily characterized by mass spectrometry. However, the relative intensities of ions detected by mass spectrometry are not always representative of the actual sample contents. SDS-PAGE can also be used to quantify conjugation yields, but the mass shift upon addition of a peptide was often poorly resolved from the non-conjugated band for smaller peptides such as Ova-I. To confirm that mass spectrometry could be used to provide reliable estimates of the conjugation efficiencies, these data were compared to densitometry measurements for SDS-PAGE analyses of MS2 PKROva-I and MS2 PKR-Eα samples. The observed conjugation levels were similar between the two techniques (SI Figure S5). Because mass spectrometry yielded better resolution than SDS-PAGE for all peptides and largely agrees with SDS-PAGE conjugation efficiency, this technique was used to determine the level of conjugation for all experiments reported herein.

To demonstrate the site-selectively of peptide conjugation to MS2, various control reactions were compared to the standard tyrosinase conjugation of tyrosine-tagged Ova-I peptide. As expected, no reaction occurred when the reaction did not contain any megaTyr enzyme (SI Figure S6). To demonstrate that the reaction is selective for cysteine residues, the cysteines on MS2 were first blocked using *N*-ethylmaleimide (NEM), then after removal of excess NEM, no reaction occurred between MS2 PKR-NEM and Ova-I using megaTyr. Lastly, two peptides that do not contain a tyrosine residue were added to MS2 PKR with megaTyr, and no reaction occurred, as predicted. These results corroborate previously published work on the site selectivity of tyrosinase conjugations between cysteine containing MS2 mutants and tyrosine-containing peptides.^29^

The degree of peptide coupling can be controlled by changing the peptide concentration. When reacted with increasing ratios of peptide to MS2 PKR, the resulting modification level increased (Figure 2c). Precise control over the degree of conjugation would be useful if multiple types of peptide antigens are needed in one capsid, or if a small molecule drug is to be used in addition to the peptides inside MS2. Constructing vaccines with multiple peptide antigens per capsid would be particularly useful for cancer vaccine applications, in which cell heterogeneity is inherent. To demonstrate the addition of multiple peptides to MS2, both Ova-I and Eα were reacted with MS2 PKR simultaneously using megaTyr (SI Figure S4). Using different ratios of each peptide in the reaction with MS2 PKR demonstrated that the modification yield of each peptide can be easily controlled. The high efficiency, specificity, and control that megaTyr allows for oxidative coupling of peptides to MS2 makes it a very useful reaction for the construction of a wide variety of peptide vaccines.

### MS2-peptide vaccine candidates remain assembled and do not contain free peptide

The three antigens that were primarily used in this study were the MHC-I antigen Ova_257-264_ (Ova-I, SIINFEKL), the MHC-II antigen Eα_52-68_ (Eα, ASFEAQGALANIAVDKA), and the MHC-II antigen Ova_323-339_ (Ova-II, ISQAVHAAHAEINEAGR), with a GGY tag added to the Cterminus of each for tyrosinase conjugation. Each peptide was conjugated to the C87 residue on the inside of MS2 PKR using megaTyr until a high labeling percentage was reached, verified by mass spectrometry (Figure 3a). When the reactions were complete, megaTyr was removed. For quick and effective removal, megaTyr was added to nickel resin before use in peptide conjugation reactions, and each reaction was filtered when complete to remove megaTyr. The resulting filtrate did not show tyrosinase activity for 24 h (SI Figure S7). Additionally, excess peptide was removed via buffer exchange, and complete removal of excess peptide was confirmed by reversed-phase high-performance liquid chromatography (RP-HPLC) (Figure 3b). The product MS2-peptide vaccine candidates maintained a high degree of capsid labeling with an indetectable amount of free peptide.

**Figure 3.**
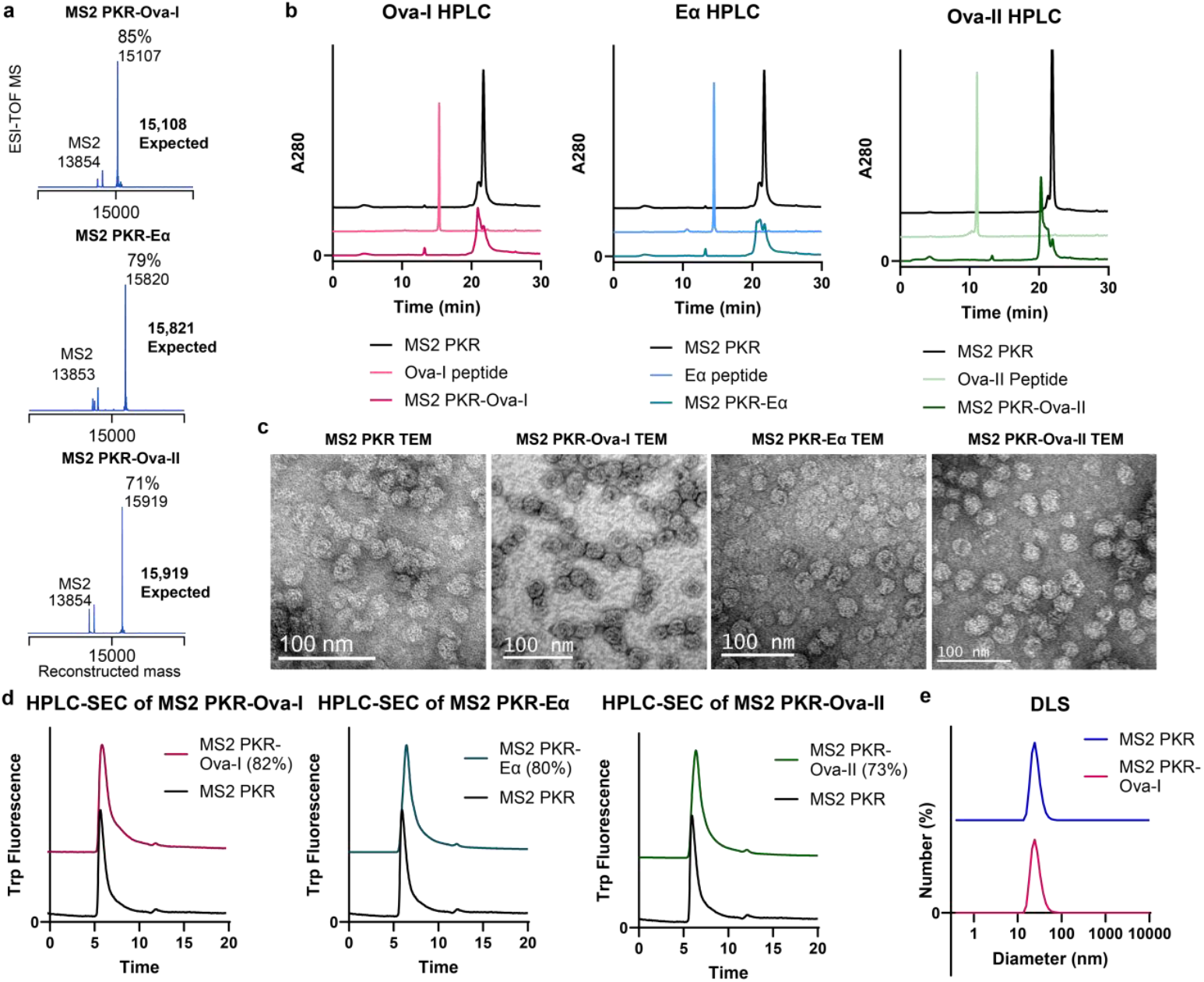
Synthesis of MS2-peptide vaccine candidates, a) ESI-QTOF spectra of MS2 PKR-Ova-I, MS2 PKR-Ea, and MS2 PKR-Ova-II conjugations, b) RP-HPLC chromatogram of MS2 PKR-peptide conjugates show complete removal of free peptide, c) TEM images of MS2 PKR capsid and MS2 PKR-peptide conjugates, d) HPLC-SEC analysis of MS2 PKR-peptide conjugates show the constructs are stable capsids. MS2 dimers and monomers, that appear at 10 min and 15-18 min respectively, are not present, e) DLS measurments of MS2 PKR and MS2 PKR-Ova-I. The MS2 PKR and MS2 PKR-Ova-I capsids measured 26.3 ± 2.0 nm and 26.7 ± 1.9 nm, respectively.

When peptide antigens were conjugated to MS2 PKR, the capsids maintained their assembly state. Transmission electron microscopy (TEM) was used to image the capsids. When labeled with a high amount of peptide, the MS2-peptide conjugates were indistinguishable from MS2 PKR alone, with no signs of disrupted capsid assembly (Figure 3c). The size ranges of all of the capsids remained comparable as well, each with a diameter close to the wild-type diameter of 27 nm (SI Figure S2). Additionally, HPLC-size exclusion chromatography (HPLC-SEC) traces of the MS2-peptide conjugates matched the chromatogram of non-conjugated MS2 PKR for all three antigens tested: Ova-I, Eα, and Ova-II (Figure 3d). Assembled MS2 capsids elute around 6-7 minutes, MS2 dimers elute around 10 min, and MS2 monomers elute between 15-18 min; only the assembled capsid was observed.^19^ Lastly, dynamic light scattering (DLS) measurements of peptide-coupled and empty capsids were the same, as expected. MS2 PKR measured 26.3 ± 2.0 nm and MS2-Ova-I was 26.7 ± 1.9 nm via DLS with N=3 biological replicates (Figure 3e). Based on these observations, we conclude that MS2-peptide vaccines were made with high conjugation, all excess peptide was removed, and the final constructs were stable, assembled capsids.

### MS2-peptide conjugates promoted activation of DC2.4s, while free peptide antigens did not

To demonstrate that MS2-peptide vaccines can activate DCs, four markers were measured using flow cytometry. Increased presentation of MHC-II, CD40, CD80, and CD86 are well-studied signs of DC activation.^21,22^ It was demonstrated in Figure 1c that MS2 capsids lacking additional antigens activate DC 2.4 cells well, making MS2 a particularly useful carrier for vaccine antigens because the capsid itself serves as an adjuvant.

Peptide antigens alone can directly load into MHC molecules that are already presented on the surface of cells *in vitro*; however, this does not activate the cells.^34^ Immature DCs can present antigens, but without activation they have low expression levels of co-stimulatory molecules such as CD40, CD80, and CD86, which are required for T-cell activation.^35^ As expected, DC2.4 cells treated with only the Ova-I and Eα peptides did not activate DC2.4 cells, so even if direct loading of the peptides into surface presented MHCs occurs, the cells will be lacking the upregulation of co-stimulatory molecules for Tcell activation (Figure 4a, 4b). Additionally, if the peptides do not make it inside of the cells to follow an MHC-I or MHC-II loading pathway, the effect of direct external MHC-loading will be short lived *in vitro* and likely would not lead to T-cell activation *in vivo* due to rapid protease degradation of free peptides.^36–38^

**Figure 4.**
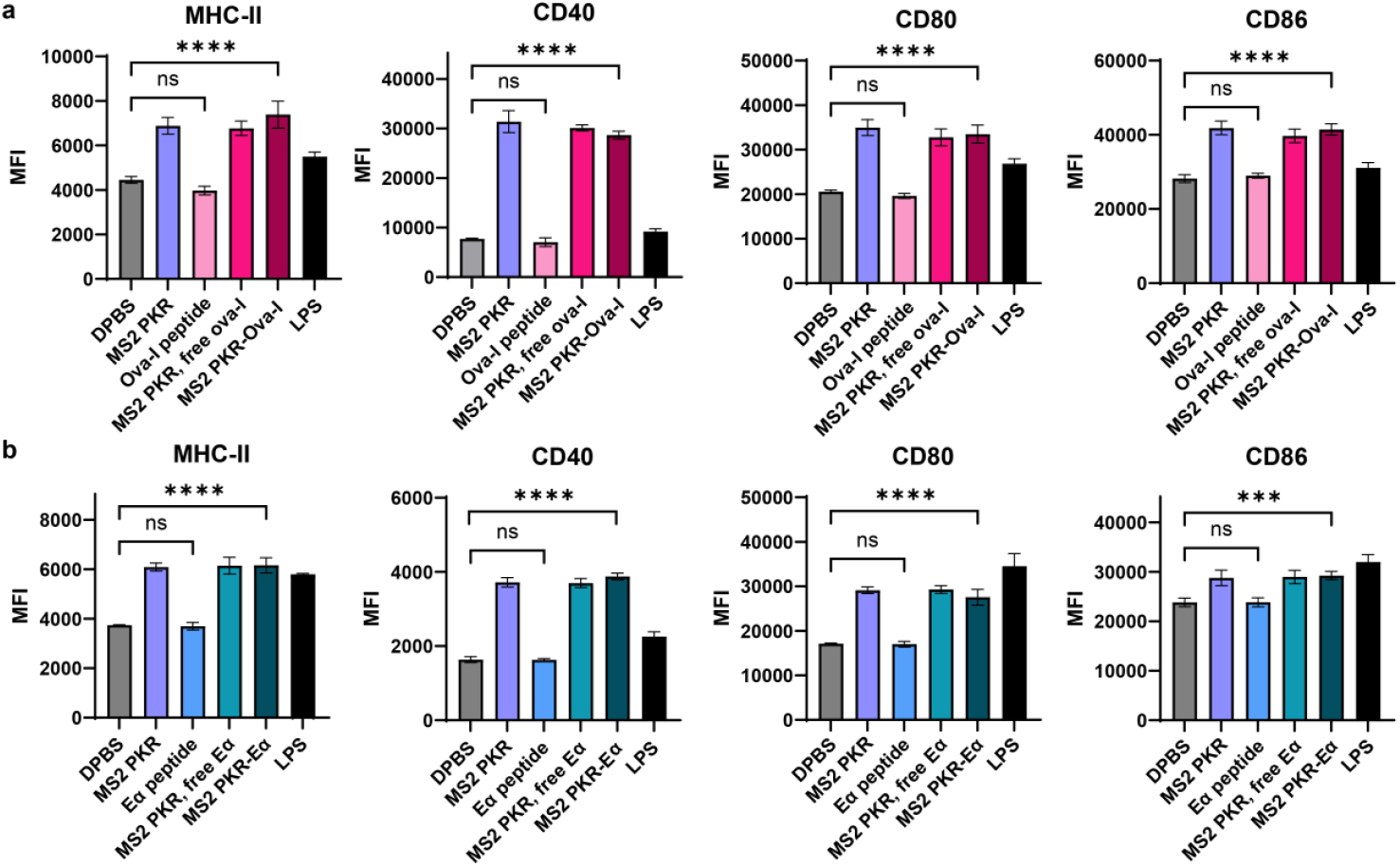
Activation of DC2.4 cells. Surface expression of MHC-ll, CD40, CD80 and CD86 on DC2.4 cells after treatment with a) MS2 PKR-Ova-I or b) MS2 PKR-Ea and controls for 24 h, analyzed by flow cytometry. LPS controls were treated with 1 ng of LPS per well. N=3 biological replicates are shown.

All samples that contained MS2 PKR had similar levels of DC activation with highly upregulated markers compared to the DPBS control and free peptides (Figure 4a, 4b). Activation of DC2.4 cells was maintained with the addition of peptide antigens either conjugated or non-conjugated to MS2. Therefore, MS2-peptide vaccine constructs maintained the adjuvant activity of MS2 PKR, which activates DCs well. This DC activation aids in promoting the processing and presentation of delivered peptide antigens while inducing the production of costimulatory molecules required for T-cell activation.

### MS2 induced prolonged cell surface display of MHCs

The mechanism by which MS2 PKR capsids are taken up by cells appears to be energy dependent and likely involves an endocytic or phagocytic mechanism.^39^ This mechanism resembles exogenous antigen uptake in cells, suggesting it is well suited for delivery of MHC-II antigens such as Eα. However, endogenous MHC-I antigens like Ova-I delivered in this way can be cross-presented by DCs.^40^ The peptide antigens that bind to MHC-I molecules are usually short in length, between 8 and 11 amino acids and are known to bind to the MHC primarily using anchor residues at their N and C-termini.^41,42^ A GGY tag was added to the C-terminus of each peptide antigen in this work to enable tyrosinase bioconjugation with MS2, which could interfere with MHC-binding. While some reports have indicated that it is possible for MHC-I peptides to bind with a Cterminal extension,^42,43^ it is possible that the GGY tag at the C-terminus of the Ova-I peptide will need to be degraded before the Ova-I peptide can bind to its H-2K^b^ MHC. Because the tyrosinase linkage between MS2 and the peptide antigens is irreversible and the MS2 capsid should protect from protease degradation (see Figure 6), any MHC presentation observed from MS2-peptide constructs should be the result of peptide delivery and presentation arising from an MHC presentation pathway.

DC2.4 cells were treated for 24 h with the MS2 PKR-Ova-I conjugates and controls, then stained with an antibody dye that responds to the H-2K^b^ MHC bound to the Ova-I peptide. Highly increased staining for MHC-Ova-I presentation was observed at all measured concentrations of MS2 PKR-Ova-I compared to the DPBS control (Figure 5a). In addition, some staining was observed for the samples treated with MS2 PKR only, even though there was no Ova-I peptide in those samples. This is likely because the antibody dye is somewhat promiscuous towards the MHC H-2K^b^ without the peptide bound (see further verification of this in the next section). Because the MS2 PKR alone has been shown to activate DCs, MS2 PKR likely induced increased surface presentation of the H-2K^b^ compared to the DPBS control. Subsequent T-cell activation needs to be measured to confirm the observed MHC staining of cells treated with only MS2 PKR is not a result of actual Ova-I presentation (*vide infra*). However, compared to samples treated with only MS2 PKR, MHC presentation was higher for MS2 PKR-Ova-I-treated samples at 200 and 1,000 nM.

**Figure 5.**
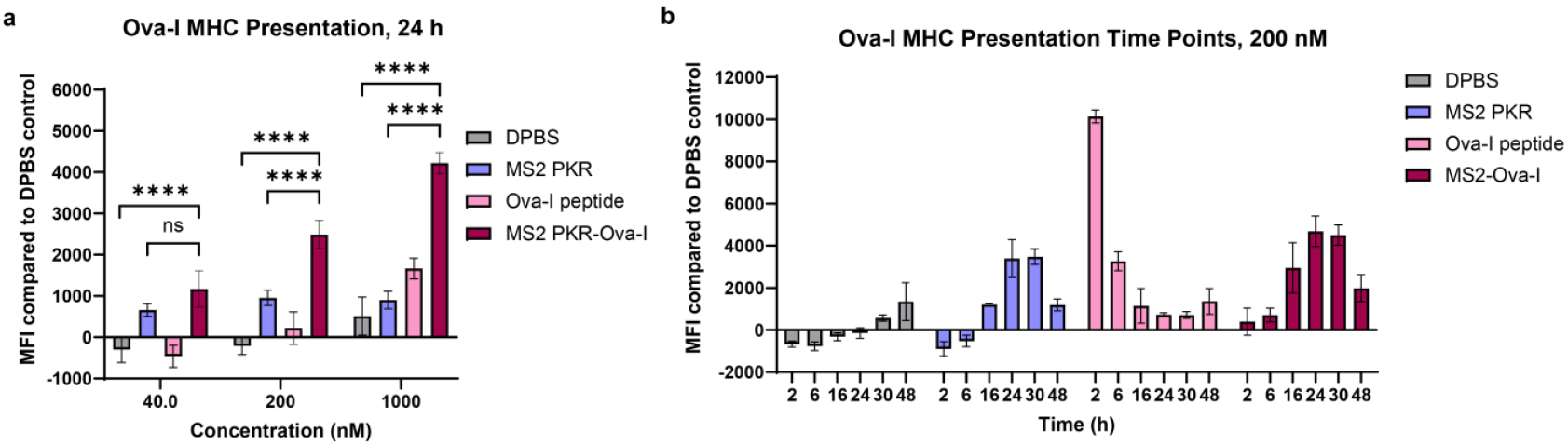
Trends in MHC-I presentation identified using flow cytometry, a) Flow cytometry analysis of Ova-I peptide presentation on MHC-I molecules of DC2.4 cells treated with MS2-Ova-I and controls at 40, 200, and 1000 nM concentrations of peptide for 24 h. b) Time points of Ova-I MHC-I presentation on DC2.4 cells after treatment with MS2-Ova-I and controls for 2, 6, 16, 24, 30, and 48 h at 200 nM of peptide, analyzed by flow cytometry. MFI values reported have the mean MFI of the DPBS controls subtracted from them. N=3 biological replicates are shown.

MHC presentation of Ova-I was also measured at 200 nM over various time points from 2 h to 48 h. An immune response curve was observed for the MS2 PKR-Ova-I sample, with peak presentation occurring around 24 to 30 h (Figure 5b). Samples that contained only the Ova-I peptide showed a sharp uptick in signal at 2 h, which quickly decreased. This immediate spike of MHC presentation is likely the result of the peptide’s ability to load directly onto already surface-presented MHCs, rather than actual cross presentation by the cells.^34^ The immediate spike in MHC-presentation disappears quickly due to protease degradation of the peptide. Samples treated with only MS2 PKR show a similar but slightly weaker immune response curve as the MS2 PKR-Ova-I samples. As mentioned previously, this likely occurred because the MS2 PKR activated the DCs, leading to increased cell surface presentation of MHC H-2K^b^ that the dye seems to bind to even without Ova-I peptide present.

To be cross-presented by DC2.4 cells, the peptide must be cleaved from MS2 PKR, but the MS2 PKR-Ova-I constructs do not contain a cleavable linker. To see if a cleavable linker would be beneficial, a valine-citrulline (val-cit, VC) cathepsin-cleavable linker was added between the peptide and the GGY tag, was conjugated to MS2 PKR and the DC2.4 cells were treated over the same series of time points (SI Figure S8). The resulting MHC presentation was very similar to the constructs without the val-cit tag, so the cleavable linker was determined to be unnecessary for MHC presentation in this case.

In summary, MHC presentation of peptides delivered using MS2 PKR cannot be confidently concluded using flow cytometry because the antibodies were observed to be promiscuous towards the MHC without the correct peptide bound.

Therefore, the monitoring of subsequent T-cell activation is required to confirm MHC presentation of MS2-delivered peptides. However, these flow cytometry experiments for MHC-presentation were still very useful to observe the immune response trends in MS2-peptide vaccines and controls and to measure peak MHC presentation times to inform the design of further experiments.

### DC2.4 cells treated with MS2-peptide conjugates activated Ova-I-specific B3Z T-cell hybridomas

To induce the adaptive immune response needed for disease protection, MS2-delivered peptide antigens need to be presented in MHC molecules by DC cells, which then must activate T-cells. B3Z T-cell hybridomas are a type of immortalized T-cell that has been engineered to res pond specifically to the Ova-I peptide only when it is bound to H-2K^b^ MHC. B3Z cells have a LacZ gene that encodes for the expression of the βgalactosidase enzyme when the cells are activated.^44,45^ This allows B3Z activation to be measured using a β-galactosidase assay in well plates.

To measure if the MS2-peptide vaccine candidates lead to MHC presentation that can stimulate B3Z cells, DC2.4 cells were first treated with MS2-peptide conjugates for 24 h, followed by co-culturing with B3Z cells for 22 h. The B3Z cells were then lysed and treated with chlorophenol red-β-D-galactopyranoside (CPRG), which absorbs at 570 nm when cleaved by β-galactosidase (Figure 6a).^44^ Peak MHC presentation from the MS2 PKR-Ova-I constructs was determined to occur around 24 h (Figure 5b), so the DC2.4 cells were co-cultured with B3Z cells after 24 h of treatment. The MS2 PKR-Ova-I construct produced the highest B3Z activation compared to the controls. The B3Z activation of MS2 PKR co-delivered with free Ova-I was a result of multiple additive effects that led to increased MHC presentation of the peptide. Most likely, the MS2 PKR activated the DC2.4 cells to present more H-2K^b^ MHCs, then the free Ova-I loaded directly into the MHCs externally. Some free peptide may also have been brought into cells alongside MS2 PKR; however, MS2 PKR and Ova-I do not form noncovalent complexes, so this is likely a minimal effect (see SI Figure S9). The B3Z activation of DC2.4 cells treated with Ova-I only is the result of the ability for Ova-I to load externally into MHC H-2K^b^s that were already surface presented. B3Z activation by Ova-I-treated DC2.4 cells decreased the longer DC2.4 cells were treated because the peptide degraded in the media over time. This is supported by B3Z activation data where DC2.4 cells were treated with each sample for only 8 h before B3Z co-culture. Higher B3Z activation was observed for Ova-I peptide with 8 h before co-culture with B3Zs, compared to 24 h, due to less of the peptide having been degraded in the media with the shorter amount of time (SI Figure S10). These data agree with the flow cytometry time points for MHC presentation (Figure 5b), which show a high spike of initial staining for Ova-I only samples before proteases degrade the peptide over time.

**Figure 6.**
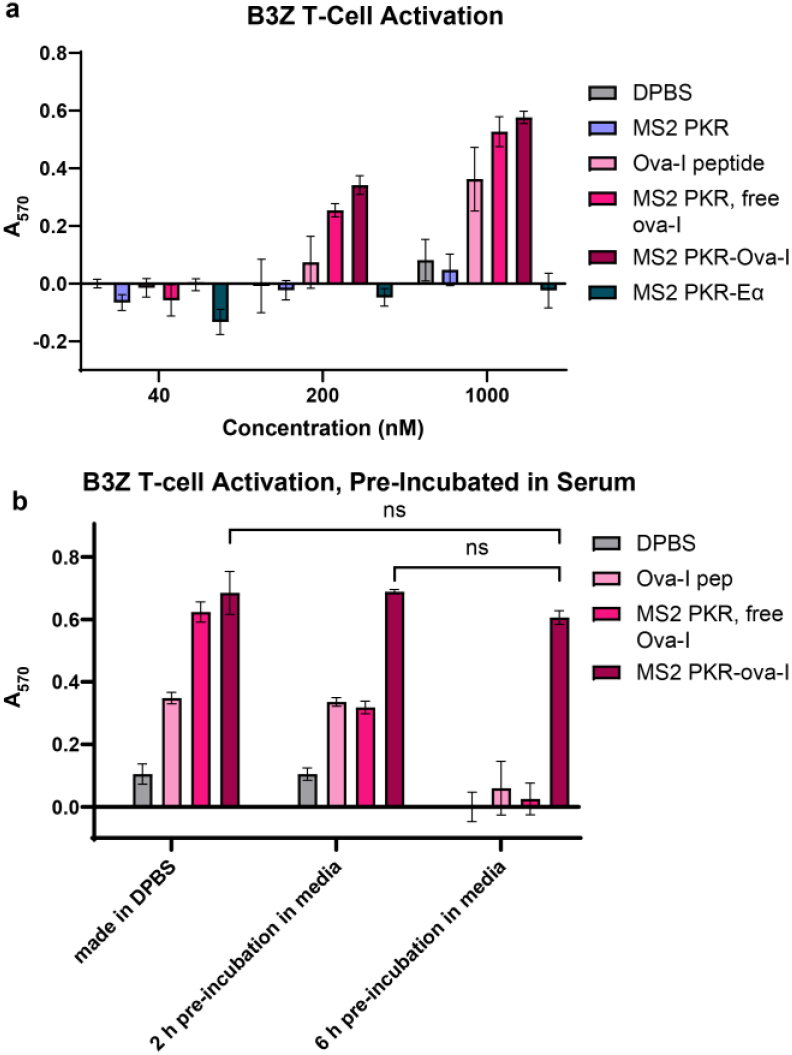
B3Z and OTHZ T-cell activation, a) DC2.4 cells were treated with MS2-Ova-I and controls for 24 h, then co-culturcd wtih B3Z T-cells, b) B3Z activation with samples that were made freshly in DPBS or prc-incubatcd with FBS-suplimented cell culture media for 2 h or 6 h at 37 °C. A_57O_ corresponds to the absorbance of CPRG after it has been cleaved by the P-galactosidase in activated B3Z cells. N=3 biological replicates are shown.

No B3Z activation was observed for the samples treated with only MS2 PKR and MS2 PKR-Eα. The lack of B3Z activation in samples treated with only MS2 PKR was notable because the flow cytometry experiments in the previous section showed staining from the antibody that was supposed to be specific to MHC H-2K^b^ bound to Ova-I even when there was no Ova-I peptide in the system. These data confirm that the increased MFI of DC2.4 cells treated with only MS2 PKR in Figure 5 was not MHC presentation of the peptide. Instead, MS2 PKR activated the DC2.4 cells to cause upregulation of MHC H-2K^b^, which the antibody bound nonspecifically without Ova-I present. Additionally, MS2 PKR conjugated to Eα did not activate B3Z cells, which demonstrated that the B3Z T-cell response was specific to only the Ova-I peptide. Taken together, these B3Z data confirm that DC2.4 cells treated with MS2 PKR cannot activate B3Z T-cells without the addition of the correct peptide.

For *in vivo* vaccination to be successful, any peptides being delivered will need to be protected from the protease degradation that rapidly occurs in the body. Our platform of peptide delivery is unique because it allows for internal loading of the peptide to the MS2 capsid, which has the potential to protect the peptide from protease degradation. Previous data in this work have shown that the Ova-I peptide was degraded by proteases in fetal bovine serum (FBS)-supplemented cell culture media over time (Figure 5b). To test if the MS2 PKR capsid protects the Ova-I peptide from protease degradation in serum, samples of MS2 PKR-Ova-I and controls were first incubated in FBS-supplemented cell culture media for 2 h and 6 h at 37 °C. Next, the pre-incubated samples were used to treat DC2.4 cells for 24 h followed by B3Z co-culture and a β -galactosidase readout for T-cell activation. Excitingly, the MS2 PKR-Ova-I samples maintained high activation of B3Z cells, with no significant change in activation level even after 6 h of exposure to FBS-supplemented cell culture media, which naturally contains proteases (Figure 6b). Both controls containing free Ova-I peptide showed substantial degradation of the peptide over time, and B3Z activation was completely depleted for both controls after 6 h of pre-incubation in protease-containing cell culture media. These data suggest that the MS2 PKR capsid provides strong protection of internally loaded peptides and indicates that this platform for peptide delivery would likely be successful *in vivo*.

## CONCLUSION

MS2 mutants have previously been established as effective delivery vehicles for small molecule drugs, with the delivered cargo reaching cytosolic targets.^46^ Here, we have expanded the application scope of MS2 by studying the use of MS2 for delivering peptide antigens for vaccination. The MS2 capsid was demonstrated to be precisely modified with a variety of peptide antigens, protect internal peptide cargo from protease degradation, and serve as an adjuvant to aid the immune response to delivered peptide antigens. MS2 PKR activated dendritic cells to upregulate markers involved in MHC presentation and T-cell activation. Additionally, peptide antigens delivered using MS2 PKR were taken up into cells and had prolonged MHC presentation. DC2.4 cells treated with MS2-delivered Ova-I antigen successfully activated antigen-restricted B3Z T-cell hybridomas. Taken together, the cell uptake, DC activation, MHC-presentation and subsequent T-cell activation of MS2-peptide antigen conjugates demonstrated that MS2-peptide vaccine candidates stimulate the adaptive immune response needed to provide protection against a target pathogen.

The faster vaccines can be built, the quicker a strong response can be launched against any new pathogens that begin spreading. MS2 provides a vaccine scaffold in which peptide antigens can be quickly and easily interchanged to target virtually any pathogen with known peptide antigens. The antigens are not limited to communicable diseases; for example, peptide antigens specific to cancer could be used make cancer vaccines that serve as immunotherapies. Additionally, multiple types of peptide antigens could be added in each capsid for stronger protection against diseases, or to prevent resistance pathways in cancer. Based on the *in vitro* data presented here, MS2-peptide vaccines show strong potential for providing immunological protection against a wide variety of pathogens with known immunodominant peptide antigens.

## METHODS

### Cloning of MS2 Variants

As described in a previous report, EMPIRIC cloning was used with MS2 coat protein entry vector plasmids that contain GFP fragments that are flanked with BsaI recognition sequences.^39^ Two single-stranded DNA primers were extended by overlap extension PCR and purified using the Wizard SV Gel and PCR Clean-Up System (Promega). Using a 1:10 ratio of insert to vector, Golden Gate cloning was used to replace entry vector GFP sequences with each desired MS2 sequence, followed by treatment with DpnI to remove template DNA. Ligated plasmids were transformed into DH10B *Escherichia coli* competent cells and grown on LB Agar plates with chloramphenicol overnight at 37 °C. Plasmid DNA was purified using the ZymoPURE Plasmid Miniprep Kit (Zymo) and then sequenced.

### Expression and Purification of MS2 Variants

MS2 plasmids were transformed into DH10B *E. coli* cells and grown overnight on LB Agar plates with chloramphenicol at 37 °C. One colony from the plate was used to grow a 10 mL overnight culture in LB with chloramphenicol at 37 °C for 16 h. A 1 L volume of 2XYT media with 25 µg/mL chloramphenicol was inoculated with the overnight culture and incubated at 37 ºC until the optical density (OD) reached 0.6 (typically about 3 h). At that point, arabinose was added to a final concentration of 0.1 % w/v to induce protein expression. Cells were harvested after 22 h of expression and lysed using sonication.

To purify MS2, the capsids were first precipitated using 50% saturated ammonium sulfate overnight at 4 °C. FPLC was used to purify the capsids further. The HiScreen Capto Core700 (Capto Core) column was used with an isocratic flow of 10 mM sodium phosphate buffer with 2 mM sodium azide at pH 7.4. Eluent from the Capto Core column was then desalted to remove residual ammonium sulfate salt using the HiPrep 26/10 Desalting column (Cytiva). MS2 capsids without KR mutations have completed purification at this step. MS2 variants with KR mutations were further purified using the HiPrep Heparin FF 16/10 column (Cytiva) with 20 mM sodium phosphate buffer, pH 7.5 and a gradient from 0 to 100 % of 20 mM sodium phosphate buffer with 2 M NaCl, pH 7.5. Eluted fractions containing MS2 were desalted into 10 mM sodium phosphate buffer with 2 mM sodium azide using the HiPrep 26/10 Desalting column and stored at 4 °C. Due to stability concerns, these samples were never frozen. The mass of each MS2 variant was confirmed by QTOF-ESI mass spectrometry.

### Expression and Purification of megaTyr tyrosinase

MegaTyr was purified and characterized as in a previous report.^30^ Briefly, a frozen stock of BL21(DE3) cells containing an expression plasmid for megaTyr was used to grow a 10 mL start culture in LB with kanamycin for 16 h at 37 °C. The starter culture was used to inoculate 1 L of 2XYT media, then expression was induced at an OD of 0.6 using 1 mM IPTG. After 12-16 h, the cells were harvested and lysed using sonication. FPLC was used to purify megaTyr using a HisTrap HP column (Cytiva) and 20 mM Tris-HCl with 500 mM NaCl, pH 7.5 and a gradient from 10 mM imidazole to 250 mM imidazole. The elution of megaTyr was then dialyzed overnight at 4 °C into PBS with 20 µM copper(II) sulfate and the correct mass was confirmed by QTOF-ESI mass spectrometry. Glycerol was added to a final concentration of 15 % v/v and aliquots were flash frozen with dry ice and acetone before storage at -80 °C.

### Mass Spectrometry

Mass spectrometry was used to confirm the molecular weights of expressed proteins and to estimate the % modification of all MS2 conjugates. For each analysis, 1 µL of a 5 µM protein solution was injected onto an Agilent 1260 series liquid chromatograph connected to an Agilent 650 LC/QTOF mass spectrometer with electrospray ionization. Charge ladder deconvolution was performed using Agilent MassHunter BioConfirm Software 10.0 and data were graphed using Chartograph software (http://chartograph.com).

### Dye Modification

A 10 mM stock solution of AlexaFluor 594 C5 maleimide was prepared in DMSO. To 50 µM of each MS2 N87C variant in 10 mM sodium phosphate buffer with 2 mM sodium azide at pH 7.4, 5 equiv of AlexaFluor 594 C5 maleimide was edded. The samples were agitated on a rotator for 2 h at 4 °C. Excess unmodified dye was removed using seven rounds of buffer exchange (5X dilution each round) through Amicon Ultra-0.5 mL 100 kDa MWCO centrifugal filters. Modification was confirmed using QTOF-ESI-MS. Each variant was modified to 75-95% by monomer, equivalent to 135 to 171 fluorophores per capsid. The concentration of MS2-fluorophore was normalized for subsequent experiments, such that the amount of dye was the same for each sample.

### Cell Uptake and Flow Cytometry

DC2.4 cells were received from the UC Berkeley Cell Culture Facility and were maintained in RPMI + 10% FBS at 37 °C with 5% CO_2_. Each well of a flat-bottom 96-well plate was seeded with 70,000 DC2.4 cells before incubation overnight at 37 °C with 5% CO_2_. Each treatment condition was prepared as a 4X concentrated stock in DPBS at pH 7.4 with the total amount of dye normalized in each sample. To treat the cells, all media was removed by aspiration from each well of the 96 well plate and replaced with 150 µL of fresh media and 50 µL of each 4X stock treatment sample. All samples were run in biological triplicate for 1 h at 37 °C with 5% CO_2_. Next, each well was washed twice with 200 µL of 20 U/mL heparin in DPBS. Then, each well was treated with 40 µL of 0.05% trypsin for 2 min and quenched with 170 µL of RPMI + 10% FBS followed by flow cytometry analysis.

Flow cytometry was conducted using an Attune NxT flow cytometer. A minimum of 10,000 cells were analyzed for each sample. Data analysis was performed using FlowJo and mean fluorescence values were reported.

### Tyrosinase Bioconjugation of Peptides to MS2

Each peptide antigen was custom synthesized by Genscript with a GGY tag at the C-terminus for tyrosinase conjugation. Peptide stocks were made at 10 mM concentrations in DMSO. In 50 mM sodium phosphate buffer pH 6.5, various equiv of peptide were added to 20 µM of an N87C-containing MS2 variant with approximately 1,000 U/L of megaTyr for 30 min at 4 °C. The degree of modification was determined using QTOF-ESI-MS.

When a conjugate was going to be used to treat mammalian cells, megaTyr was used on-resin to allow its removal after the reaction was complete. This was important to avoid color changes in the culture media that were indicative of potential cell toxicity. The immobilized enzyme was prepared as follows: A 50% slurry of HisPur Ni-NTA resin (Thermo Fisher) was washed twice by adding 100 mM sodium phosphate buffer with 10 mM imidazole and 300 mM NaCl, pH 8.0. Each wash was followed by centrifugation at 13,000 rcf for 1 min and the supernatant was removed. To bind megaTyr to the resin, a 1:1 v/v ratio of megaTyr at 1,000 U/L to washed HisPur Ni-NTA resin was rotated in 100 mM sodium phosphate buffer with 10 mM imidazole and 300 mM NaCl, pH 8.0, for at least 1 h at 4 °C. The megaTyr-NiNTA resin was then washed twice with 100 mM sodium phosphate buffer containing 10 mM imidazole and 300 mM NaCl, pH 8.0, on a 2.0 mL Costar 0.22 µm cellulose acetate filter, then washed once more with 100 mM sodium phosphate buffer, pH 8.0. Care was taken to minimize the amount of time the resin was left dry each centrifugation step on the filter. The washed megaTyr-NiNTA resin was then resuspended to a 50% slurry in 100 mM sodium phosphate buffer, pH 8.0 and used immediately. To use the megaTyr-NiNTA resin for a peptide coupling reaction with MS2, the 50% slurry of megaTyr-NiNTA resin was added as a 1:5 dilution into a solution of 20 µM MS2 with 300 µM of peptide antigen in 50 mM sodium phosphate buffer, pH 8.0 and rotated for 1.5 h at 4 °C. The reaction must be maintained at a pH no lower than 8.0 for the megaTyr to remain on the resin. MegaTyr-NiNTa resin was removed from the reaction via filtration of the reaction through a 2.0 mL Costar 0.22 µm cellulose acetate filter.

Removal of megaTyr was confirmed via an activity assay where 20 µL of the filtered reaction was added to 0.25 mM L-tyrosine in 20 mM sodium phosphate buffer pH 6.5. Absorbance at 475 nM, corresponding to tyrosine converted to *ortho*-quinone and its subsequent products, was measured periodically over 24 h.^47^ Only samples showing no residual megaTyr activity were used in cell culture experiments. Excess peptide was removed using seven rounds of buffer exchange (5X dilution each round) through an Amicon Ultra-0.5 mL 100 kDa MWCO centrifugal filter and removal of excess peptide was confirmed using RP-HPLC. The extent of peptide conjugation to MS2 was measured using QTOF-ESI-MS and SDSPAGE. Each MS2 capsid sample was confirmed to be modified on 75-85% of its monomers, corresponding to 135 to 153 peptides per assembled capsid. The concentrations of each MS2peptide sample were normalized for all subsequent experiments.

### Reversed-Phase HPLC to Confirm Excess Peptide Removal

To confirm removal of excess peptide in MS2-peptide samples, an Agilent Technologies 1260 Infinity analytical HPLC was used with a Phenomenex C18 silica semiprep column, and a 40 min gradient of 5-95% milli-Q water with 0.1% trifluoroacetic acid (TFA) to acetonitrile with 0.1% TFA was used to confirm excess peptide was removed from MS2-peptide samples. Samples containing 20 µM of MS2 PKR, 20 µM MS2 PKR-peptide, and 300 µM of peptide (the concentration of peptide if it was not removed from the conjugation reaction) in DPBS were injected (20 µL injections) onto the column and absorbance at 280 nm was measured.

To determine if free peptide adheres to MS2 PKR, MS2 PKR was first buffer exchanged three times into DPBS using an Amicon Ultra-0.5 mL 100 kDa MWCO centrifugal filter. To 50 µM MS2 PKR in DPBS, 40 µM peptide (to match the peptide concentration of an 80% conjugated sample) was added and the sample was rotated at 4 °C for 1 h. The product was buffer exchanged two times into DPBS using an Amicon Ultra-0.5 mL 100 kDa MWCO centrifugal filter. Each sample was injected onto the HPLC with the same column and method as used for confirmation of peptide removal.

### HPLC-SEC

MS2-peptide samples were analyzed using HPLC-SEC to confirm the capsids maintained their assembly after peptide conjugation. Samples with 20 µM MS2-peptide product were injected onto an Agilent Technologies 1260 infinity HPLC with an Agilent Bio SEC-5 column using isocratic flow of 10 mM sodium phosphate buffer with 2 mM sodium azide at pH 7.4. Tryptophan fluorescence was measured with 280 nm excitation and 350 nm emission, and elution time was used to confirm the assembly state of the capsids.

### Transmission Electron Microscopy

Formvar/Carbon-coated Copper 400 mesh grids (Electron Microscopy Sciences) were first subjected to glow discharge. Subsequently, 5 µL of MS2 samples at concentrations of ∼ 50 µM were pipetted onto the grids and incubated for 2 min. The grids were then washed three times with MilliQ water. Excess liquid was removed from the grids using a Whatman filter paper. Next, 5 µL of 1% uranyl acetate (wt/vol.) solution was pipetted onto the grids and incubated for 2 min. Excess solution was removed using a Whatman filter paper, and the grids were dried under ambient conditions. The grids were then imaged with a Tecnai 12 Transmission Electron Microscope, equipped with a 2k x 2k CCD camera. Images were acquired at 23000x, 30000x, and 49000x magnification. All TEM measurements were performed at the Electron Microscope Lab at UC Berkeley.

### Dynamic Light Scattering

Each sample was concentrated to at least 50 µM, then filtered through a 2.0 mL Costar 0.22 µm cellulose acetate filter before transfer into a cuvette for measurement. Samples were measured on a Malvern Zetasizer Nano-ZS with three biological replicates. Number distribution was used to determine particle size.

### DC Activation

To each well of a 96-well flat-bottom plate, 35,000 DC2.4 cells were plated and incubated overnight at 37 °C with 5% CO_2_. To treat cells, all media was aspirated from each well and replaced with 150 µL of fresh media followed by the addition of 50 µL of 4X concentrated solutions of each treatment condition in DPBS, in biological triplicate. Each peptide-containing sample was treated with 1 µM final concentration of peptide, and MS2-containing samples were normalized to the same concentration of MS2 used in MS2-peptide samples. LPS positive control wells were treated at 5 ng/mL. Cells were treated for 24 h at 37 °C with 5% CO_2_ until harvested and stained for flow cytometry. DC 2.4 cells were used at passage 10 or below for all experiments.

The cells were prepared for flow cytometry by first gently washing each well with 200 µL of DPBS. Next, each well was treated with 100 µL of the Gibco enzyme-free cell dissociation buffer (Cell Dissociation Buffer, enzyme-free, PBS, Cat # 13151014) for 5 min at room temperature before a quench with 100 µL of RPMI with 10% FBS. Each well was mixed by pipetting up and down a couple of times to help remove adhered cells, then the supernatant was transferred to a V-bottom 96well plate. The plate was centrifuged at 400 rcf for 5 min, then washed with 200 µL of DPBS and centrifuged again. The cells were then stained with 100 µL of a 1:100 dilution of each antibody stain in DPBS with 10% FBS and 1% w/v sodium azide for 20 min at 4 °C, protected from light. Stained cells were centrifuged and washed with 200 µL of DPBS per well, then fixed with 100 µL of a 4% paraformaldehyde solution for 15-20 min. Fixed cells were centrifuged then resuspended DPBS for analysis by flow cytometry. DC2.4 cells were used at passage 10 or below for all experiments.

### MHC Loading

In each well of a flat-bottom 96-well plate, 35,000 DC2.4 cells were plated and incubated overnight at 37 °C with 5% CO_2_. Concentrated solutions (4X) of each treatment were prepared in DPBS. All media was aspirated from each well and replaced with 150 µL of fresh media and 50 µL of each corresponding 4X treatment, in biological triplicate. Treated cells were incubated for 24 h at 37 °C with 5% CO_2_ until prepared for flow cytometry.

The treated DC2.4 cells were prepared for flow cytometry as described in the DC Activation method section. Briefly, each well was washed with 200 µL of DPBS, treated with 100 µL of the Gibco enzyme-free cell dissociation buffer for 5 min, and quenched with 100 µL of RPMI with 10% FBS. The cells were transferred to a V-bottom 96-well plate and washed with 200 µL of DPBS before staining with 100 µL of a 1:100 dilution of the BioLegend PE/Dazzle™ 594 anti-mouse H-2K^b^ bound to SIINFEKL Antibody in DPBS with 10% FBS and 1% w/v sodium azide for 20 min at 4 °C, then washed with 200 µL of DPBS. The cells were fixed with 100 µL of a 4% paraformaldehyde solution for 15-20 min, then suspended in DPBS for analysis by flow cytometry. DC2.4 cells were all used at passage 10 or below.

### MHC-Loading Time Points

Samples of 20,000 DC2.4 cells were plated in each well of a flat-bottom 96 well plate and incubated overnight at 37 °C with 5% CO_2_. Each time point was measured in reverse chronological order, starting with the 30 h time point and ending with the 2 h time point, so all cells were harvested and stained for flow cytometry together. At the 30 h time point, media was removed from all wells corresponding to all time points and 150 µL of fresh media was added to each well. At each time point, 50 µL of each 4X concentrated sample (made in DPBS) was added to their corresponding wells, in biological triplicate. The plates were incubated at 37 °C with 5% CO_2_ between each time point and until harvested for flow cytometry. All DC 2.4 cells used for these experiments were passage 10 or below.

### Flow Cytometry for DC Activation and MHC-Loading

All samples for DC activation, MHC loading, and MHC loading time points were prepared for flow cytometry using these methods. Each flow cytometry stain with prepared as a 1:100 dilution in DPBS with 10% FBS and 1% w/v sodium azide at 4 °C and used immediately. To detach cells, first, all media was aspirated, and each well was washed with 200 µL of DPBS and aspirated. Next, 100 µL of Gibco Cell Dissociation Buffer Enzyme-Free PBS-based was added to each well at room temperature for 5 min, then 100 µL of RPMI + 10% FBS was added to each well. Each sample was pipetted up and down a few times to help cells detach, then transferred to a 96-well V-bottom plate. Plates were centrifuged at 400 rcf for 5 min and the supernatant was rapidly decanted via a quick upside-down flick of the plate and blotting onto a paper towel. Cells were resuspended in 100 µL of each diluted stain solution and incubated at 4 °C for 20 min, protected from light. Plates were centrifuged at 400 rcf for 5 min and the supernatant was removed by flicking. Each well was washed by resuspension in 200 µL of prechilled DPBS at 4 °C and centrifuged at 400 rcf for 5 min. Supernatant was removed and each well was fixed with 100 µL of pre-chilled 4% paraformaldehyde at 4 °C for 20 min, protected from light. Each plate was centrifuged at 400 rcf for 5 min, then supernatant was removed, and the cells were resuspended in 200 µL of pre-chilled DPBS at 4 °C. Each plate was kept on ice and protected from light until injected into the flow cytometer.

An Attune NxT flow cytometer was used to measure all samples. FlowJo was used to analyze these data and mean fluorescence values (MFI) were reported. MFI of all DPBS control samples per experiment was set as zero for all MHC-loading values reported.

### B3Z Co-culture

DC2.4 cells were received from the UC Berkeley Cell Culture Facility and were cultured in RPMI + 10% FBS at 37 °C with 5% CO_2_. B3Z T-cell hybridomas were received from the UC Berkeley Cell Culture Facility and cultured in RPMI + 10 % FBS + 1% sodium pyruvate + 0.1 % 2-mercaptoethanol at 37 °C with 5% CO_2_. All cells were maintained at passage 10 or below.

DC2.4 cells were plated at 20,000 cells per well in a U-bottom 96-well plate and incubated at 37 °C with 5% CO_2_ overnight. To treat the DC2.4 cells, all media was gently removed with a pipette and replaced with 150 µL of fresh media. Each sample was then treated with 50 µL of 4X concentrated samples (diluted in DPBS) then incubated at 37 °C with 5% CO_2_ for 24 h. After the treatment time was completed, all media was removed from each well via gentle pipetting and 100,000 B3Z T-cell hybridomas in 200 µL of media were added to each well to co-culture the treated DC2.4s with B3Z cells. Cells were cocultured for 22 h at 37 °C with 5% CO_2_.

After co-culture, a β-galactosidase assay was used measure how much β-galactosidase was produced by the activated B3Z cells. A previously reported protocol was adapted for use with our experiments.^44^ Each plate was centrifuged at 400 rcf for 5 min and supernatant was removed by gentle pipetting. Cells were washed by resuspension in 200 µL of DPBS, followed by centrifugation at 400 rcf for 5 min and gentle removal of supernatant via pipette. Each well was resuspended in 100 µL of 0.15 mM CPRG in 0.5% NP-40 in PBS and incubated at 37 °C for 18 h, protected from light. An Agilent BioTek Synergy H1 plate reader was used to shake each plate for 20 s and then measure the absorbance of each well at 570 nm, corresponding to the absorbance of the cleavage product of CPRG. The average A_570_ of all DPBS-treated control wells was set to zero for all reported values.

## ASSOCIATED CONTENT

### Supporting Information

Materials, equipment, protein and peptide sequences and properties, and supplementary figures. The Supporting Information is available free of charge on the ACS Publications website.

### Author Contributions

The manuscript was written through contributions of all authors. All authors have given approval to the final version of the manuscript.

### Notes

M.B.F. is a co-founder of and scientific advisor to Catena Biosciences, which is commercializing applications of tyrosinase-based oxidative coupling chemistry.

## ACKNOWLEDGMENT

We would like to thank the staff at the University of California Berkeley Electron Microscope laboratory for their advice and assistance in electron microscopy sample preparation and data collection.

## FUNDING SOURCES

This material is based upon work supported by the National Science Foundation Graduate Research Fellowship Program under Grant No. DGE 2146752 for HSM. Any opinions, findings, and conclusions or recommendations expressed in this material are those of the author(s) and do not necessarily reflect the views of the National Science Foundation. Additionally, the Panattoni Family Fund is acknowledged for generous financial support.

## CONFLICT OF INTEREST

MBF has a financial interest in Catena Bio-sciences, which has licensed the technology used to make the conjugates described herein, and both he and the company may benefit from commercialization of the results of this research.

## For Table of Contents only

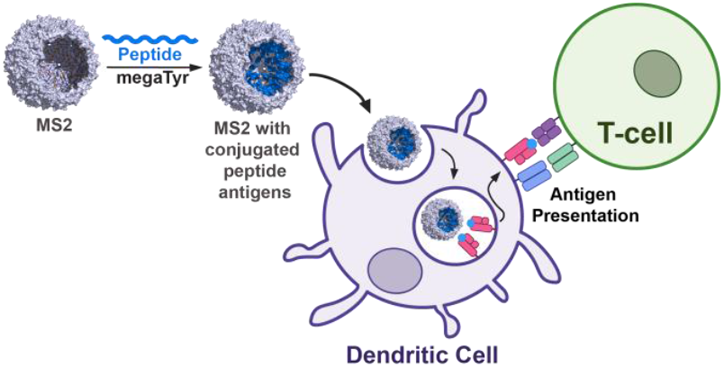

## Notes

### Summary of Updates

This version of the manuscript has been revised to add further studies demonstrating site-selectivity of the MS2-peptide conjugation, to compare conjugation efficiencies measured using SDS-PAGE and MS, and to add DLS measurements of the constructs. Additionally, further comments and explanations have been given to better introduce and explain important immunological concepts involved in this work.

