## Supporting Information for "Engineered MS2 Virus Capsids for Cellular Display of Peptide Antigens"

**Table of Contents**

**Materials……………………………………………………………………………………………………2**

**Equipment…………………………………………………………………………………………………….2**

**Supplementary Figures……………………………………………………………………………………3-12**

### Materials

#### General Reagents

DNA primers were purchased from Integrated DNA Technologies. All golden gate assembly enzymes and components were purchased from New England BioLabs and the Wizard® SV Gel and PCR CleanUp System was purchased from Promega. The ZymoPURE™ Plasmid miniprep kit was purchased from Zymo Research. DNA sequencing was performed by Genewiz from Azenta. AlexaFluor 594 C5 maleimide was purchased from Invitrogen. All peptide antigens were custom synthesized by Genscript (sequences in Figure S1).

#### Cell Lines and Cell Culture Reagents

DC2.4 cells were received from the UC Berkeley Cell Culture Facility and were cultured in RPMI + 10% FBS at 37 °C with 5% $CO_2$. B3Z T-cell hybridomas were received from the UC Berkeley Cell Culture Facility and cultured in RPMI + 10 % FBS + 1% sodium pyruvate + 0.1 % 2-mercaptoethanol at 37 °C with 5% $CO_2$. All cell lines used in experiments were at passage number 10 or below.

RPMI, 100X sodium pyruvate, 1000X 2-mercaptoethanol, and enzyme-free cell dissociation buffer (PBS-based) were purchased from Gibco. All antibody-dye conjugates used for flow cytometry were purchased from BioLegend (PE/Dazzle™ 594 anti-mouse H-2$K^b$ bound to SIINFEKL Antibody, Pacific Blue™ anti-mouse CD40 Antibody, PerCP/Cyanine5.5 anti-mouse CD40 Antibody, PerCP/Cyanine5.5 anti-mouse I-A/I-E Antibody, PE anti-mouse CD80 Antibody, Pacific Blue™ anti-mouse CD86 Antibody) with the exception of the Ea52-68 peptide bound to I-Ab Monoclonal Antibody with FITC that was purchased from Thermo Fisher Scientific.

### Equipment

An AKTA Go FPLC was used for protein purification. An Agilent 6530 LC/QTOF mass spectrometer was used for characterization of proteins. An Agilent 1260 Infinity HPLC was used for analysis of MS2-peptide conjugates. A Malvern Zetasizer Nano-ZS was used for DLS measurements. All flow cytometry experiments were performed using an Attune NxT Flow Cytometer from Thermo Fisher. All other equipment, including column names, is specified in the methods section.

**Supplementary Figures:**

**Peptide Antigens**

| Name | Abbreviation | Sequence | MHC Type | Molecular Weight | Theoretical PI |
|---|---|---|---|---|---|
| $OVA_{257\text{-}264}$ | **Ova-I** | SIINFEKLGGY | I | 1240.4 | 5.72 |
| $E\alpha_{52\text{-}68}$ | **Eα** | ASFEAQGALANIAVDKAGGY | II | 1953.1 | 4.37 |
| $OVA_{323\text{-}339}$ | **Ova-II** | ISQAVHAAHAEINEAGRGGY | II | 2051.21 | 6.00 |
| $EphA2_{883\text{-}891}$ | **EphA2** | TLADFDPRVGGY | I | 1310.4 | 4.21 |
| $hTERT_{613\text{-}626}$ | **hTERT** | EARPALLTSRLRFIPKGGY | II | 2145.5 | 10.9 |
| $TRP1_{113\text{-}126}$ | **Trp1** | CRPGWRGAACNQKIGGY (disulfide) | II | 1837.1 | 9.5 |
| Val-Cit $OVA_{257\text{-}264}$ | **VC-Ova** | SIINFEKLV{Cit}GGY | I | 1514.5 | 5.72 |

**Proteins**

| Protein | Abbreviation | Sequence | Molecular Weight |
|---|---|---|---|
| MS2 N87C | **MS2 wt** | ASNFTQFVLVDNGGTGDVTVAPSNFANGVAEWISSNSRSQAYKVTCSVRQSSA QNRKYTIKVEVPKVATQTVGGVELPVAAWRSYL**C**MELTIPIFATNSDCELIVKAM QGLLKDGNPIPSAIAANSGIY | 13,718 |
| MS2 T71K G73R N87C | **MS2 KR** | ASNFTQFVLVDNGGTGDVTVAPSNFANGVAEWISSNSRSQAYKVTCSVRQSSA QNRKYTIKVEVPKVATQ**K**V**R**GVELPVAAWRSYL**C**MELTIPIFATNSDCELIVKAM QGLLKDGNPIPSAIAANSGIY | 13,844 |
| MS2 S37P T71K G73R N87C | **MS2 PKR** | ASNFTQFVLVDNGGTGDVTVAPSNFANGVAEWISSN**P**RSQAYKVTCSVRQSSA QNRKYTIKVEVPKVATQ**K**V**R**GVELPVAAWRSYL**C**MELTIPIFATNSDCELIVKAM QGLLKDGNPIPSAIAANSGIY | 13,854 |
| Wild-type megaTyr (tyrosinase from *Bacillus megaterium*) | **megaTyr** | SNKYRVRKNVLHLTDTEKRDFVRTVLILKEKGIYDRYIAWHGAAGKFHTPPGSD RNAAHMSSAFLPWHREYLLRFERDLQSINPEVTLPYWEWETDAQMQDPSQS QIWSADFMGGNGNPIKDFIVDTGPFAAGRWTTIDEQGNPSGGLKRNFGATKEA PTLPTRDDVLNALKITQYDTPPWDMTSQNSFRNQLEGFINGPQLHNRVHRWV GGQMGVVPTAPNDPVFFLHHANVDRIWAVWQIIHRNQNYQPMKNGPFGQNFR DPMYPWNTTPEDVMNHRKLGYVYDIELRKSKRSSLE | 34,520.8 |

***Figure S1.*** Peptide and protein sequences and properties. Molecular weights and pIs were computed using the Expasy Compute pI/Mw tool. PI for citrulline (cit) was calculated using Q in place of cit in the sequence.

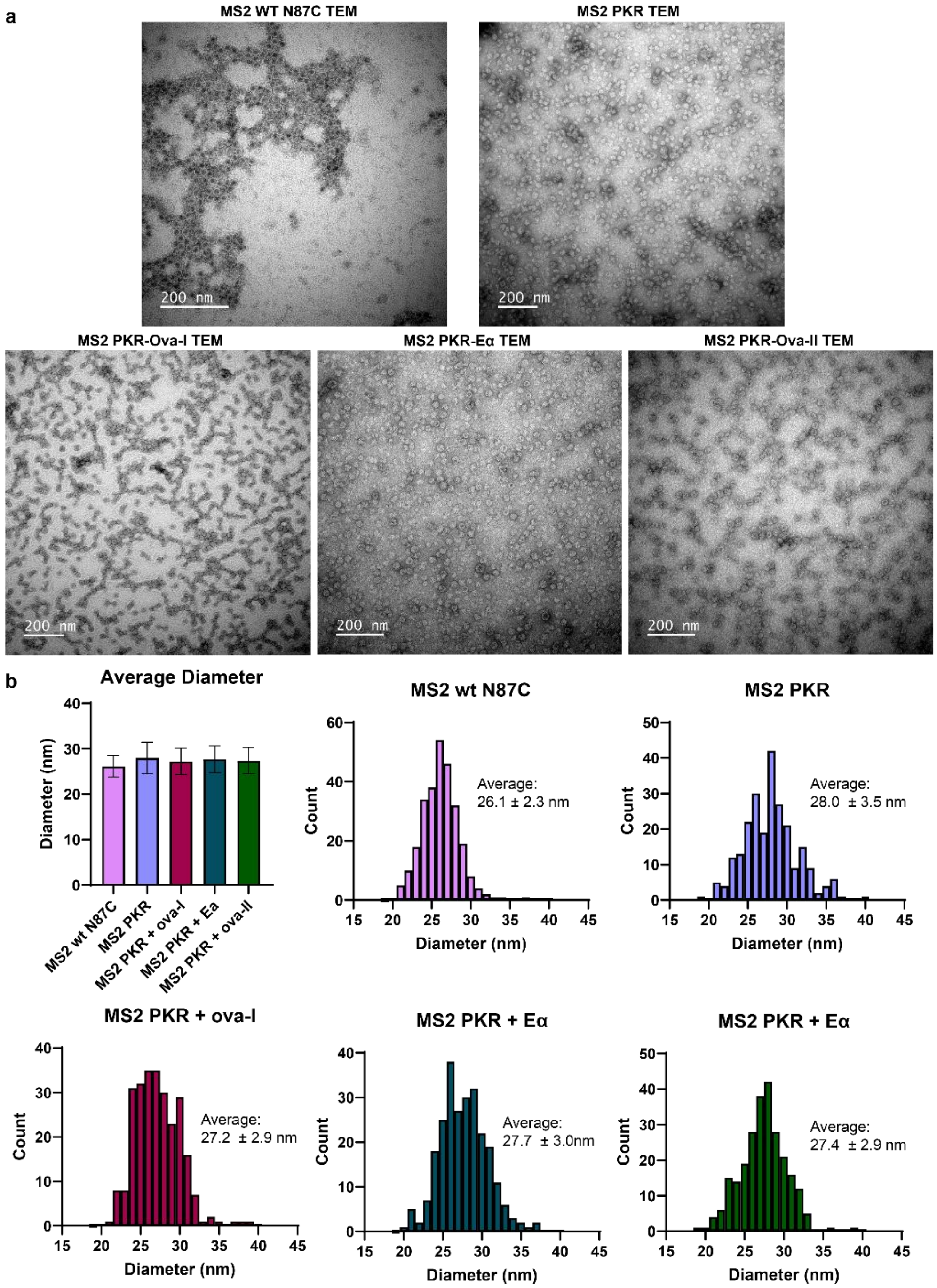


***Figure S2.*** TEM of MS2 capsids and conjugates with measurements. a) TEM images used for measurments. b) Mean capsid diameter values and histrgrams of the distribution of diameters in each sample.

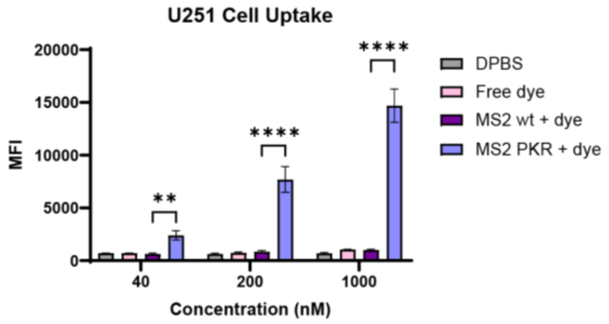


***Figure S3.*** Cell uptake in U251MG cells of Alexa Fluor 594-lableled MS2 mutants.

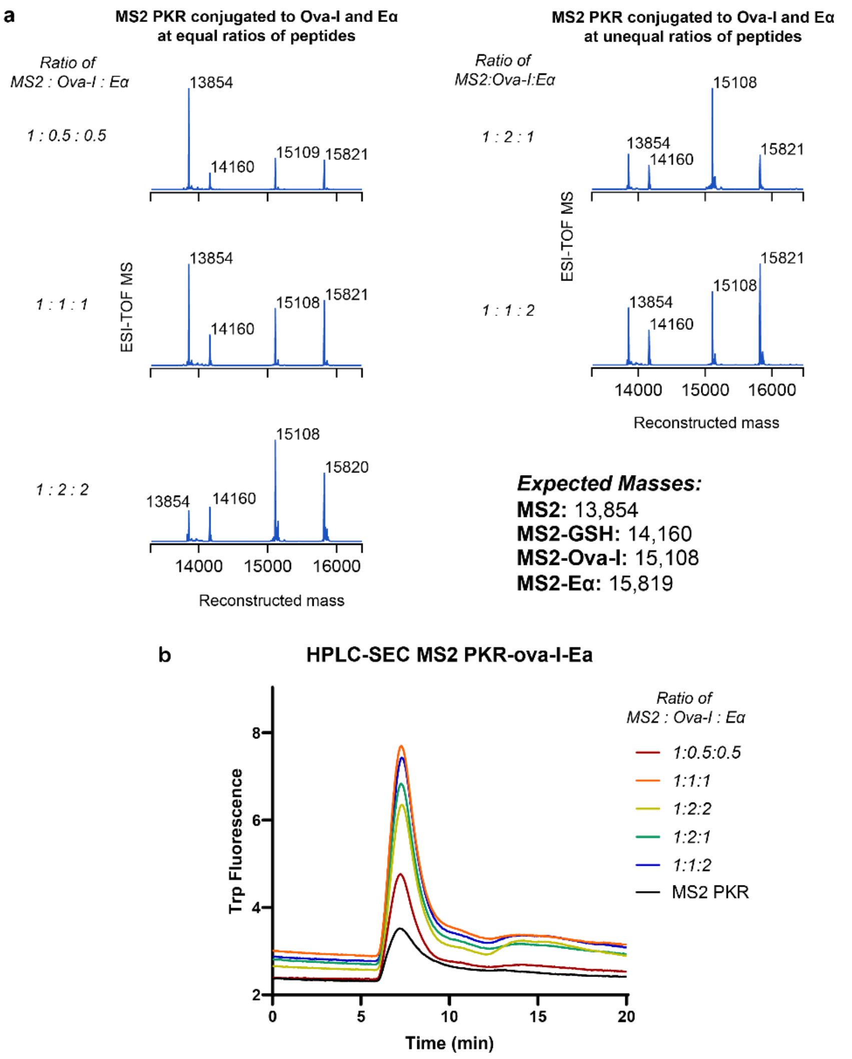


***Figure S4.*** Conjugation of multiple peptides to MS2 PKR. a) ESI-TOF-MS spectra of two peptides (Ova-I and Eα) conjugated to MS2 PKR using different ratios of peptides, noted in italics. Expected mass values are given in the figure. MS2 proteins are frequently observed as glutathione adducts, which are present in the starting protein before any oxidative coupling chemistry. These species do not participate in the coupling reactions b) HPLC-SEC chromatograms of MS2 PKR and double peptide conjugates. MS2 capsids were observed to be stable and assembled.

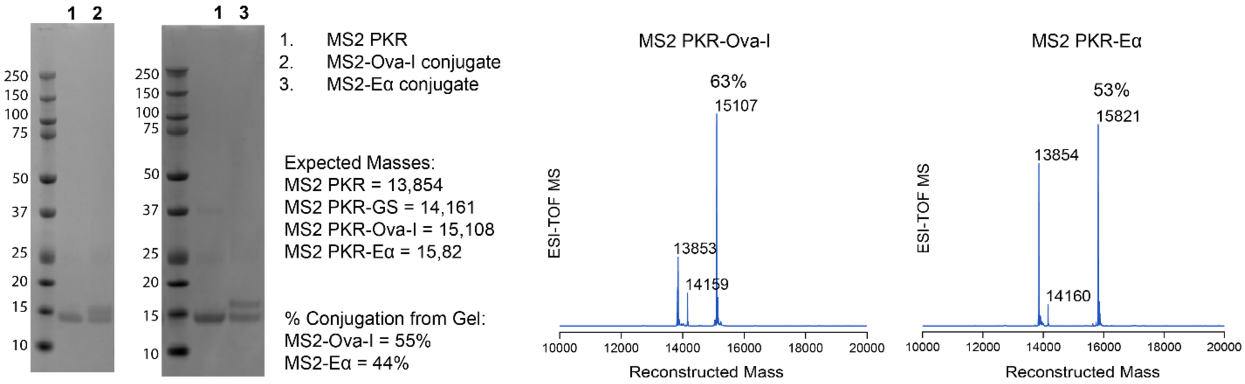


***Figure S5.*** Comparison of conjugation efficiency using SDS-PAGE and mass spectrometry. SDS-PAGE gel bands were integrated using ImageJ software. Mass spectra were integrated using Chartograph software (Chartograph.com).

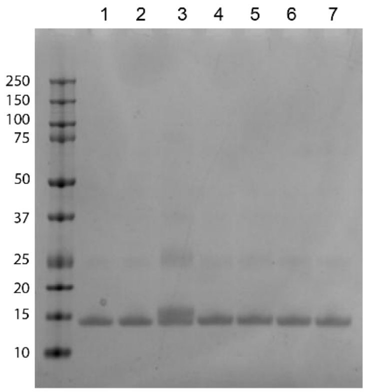


1. MS2 PKR
2. MS2 + Ova-I, no megaTYr
3. MS2-Ova-I conjugate
4. MS2-NEM conjugate
5. MS2-NEM + Ova-I
6. MS2 + K6 peptide
7. MS2 + LPS peptide

Peptide sequences:
Ova = SIINFEKLGG**Y**
K6 = WWKKWKKWW
LPS = GWKRKRFG

Expected Masses:
MS2 PKR = 13,854
MS2 PKR-Ova-I = 15,108

***Figure S6.*** Site-specificity of peptide conjugations with MS2. SDS-PAGE gel of MS2 PKR-Ova-I conjugate and controls. All peptide conjugation reactions contain megaTyr unless otherwise stated.

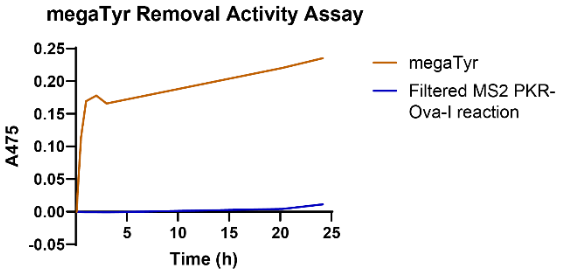


***Figure S7.*** Removal of megaTyr from MS2-peptide samples for cell studies. Samples were mixed with tyrosine, which absorbs at 475 nm when oxidized into an *ortho*-quinone using megaTyr. The megaTyr control is tested at the concentration of megaTyr in the reactions if the megaTyr was not removed. Filtered flowthrough of a conjugation of MS2 PKR-Ova-I that used megaTyr-NiNTA resin shows that megaTyr was successfuly removed.

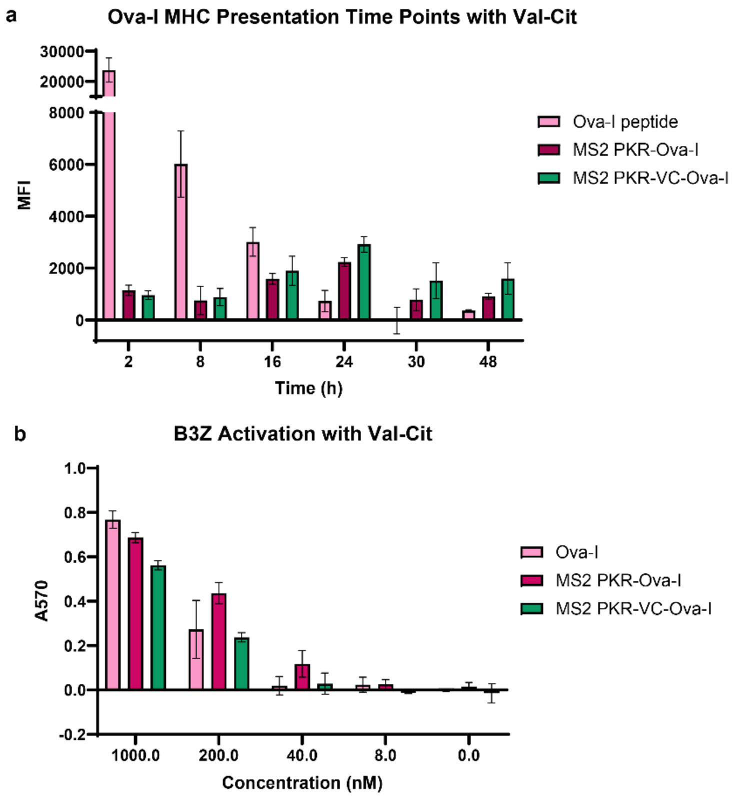


***Figure S8.*** Peptide delivery with and without a Val-Cit linker. a) MHC-loading time points of free Ova-I peptide compared to MS2 PKR-Ova-I and MHC PKR-Val-Cit(VC)-Ova-I. Mean fluorescence intensity values from flow cytometry are reported. b) B3Z activation levels with an 8 h treatment of DC2.4 cells before co-culturing with B3Z cells using ova peptide, MS2 PKR-Ova-I, and MHC PKR-VC-Ova-I, each with a peptide concentration of 200 nM. Absorbance at 570 nm corresponds to the cleavage product of CPRG.

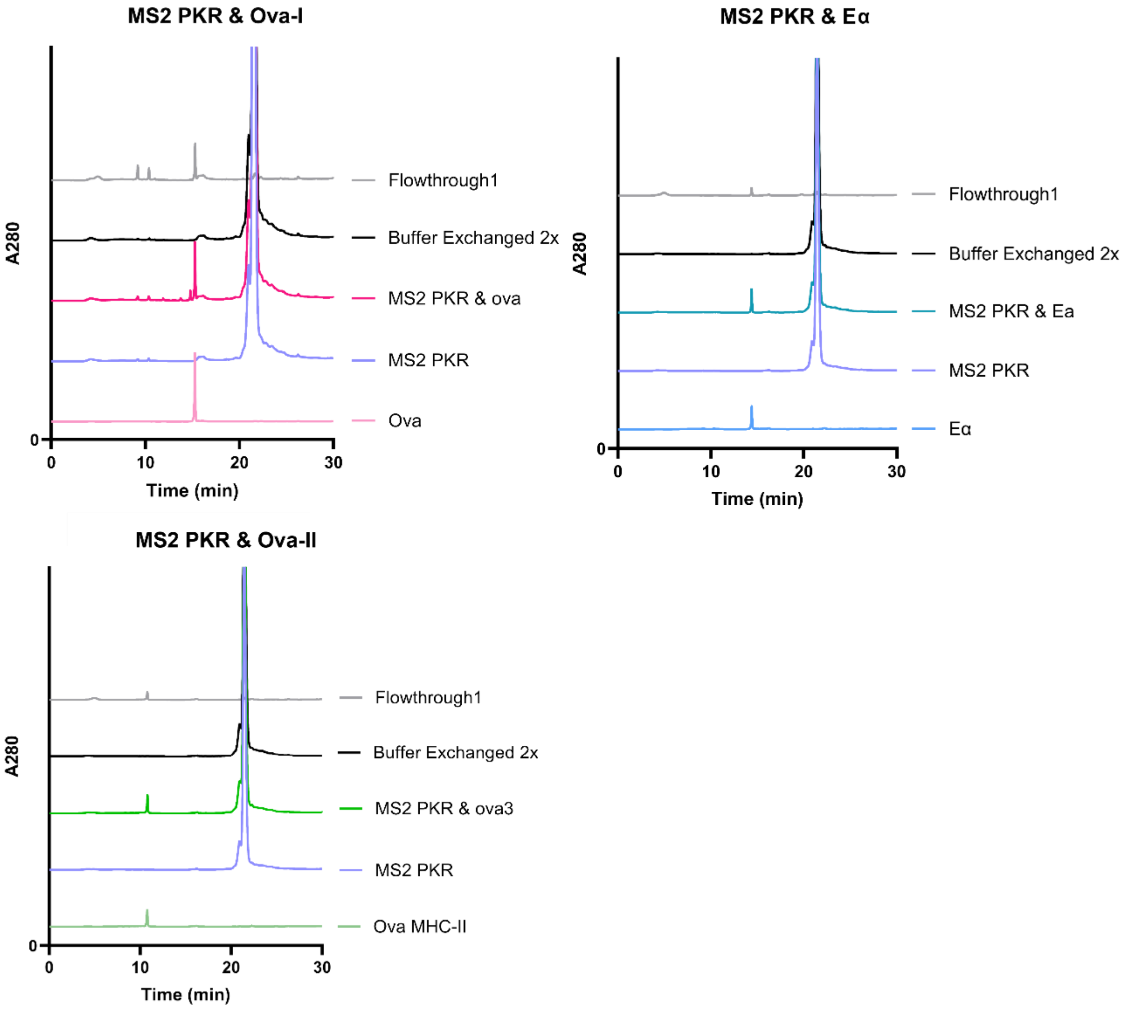


***Figure S9.*** RP-HPLC of free peptides incubated with MS2 PKR with no covalent attschement. Reverse phase HPLC of each peptide, MS2 PKR, MS2 PKR mixed with the peptide before buffer exchange (MS2 PKR & peptide), MS2 PKR mixed with the peptide that has been buffer exchanged into DPBS twice (Buffer Exchanged 2x), and the flowthrough from concentrating the MS2 PKR & peptide sample before the first round of buffer exchange (Flowthrough 1).

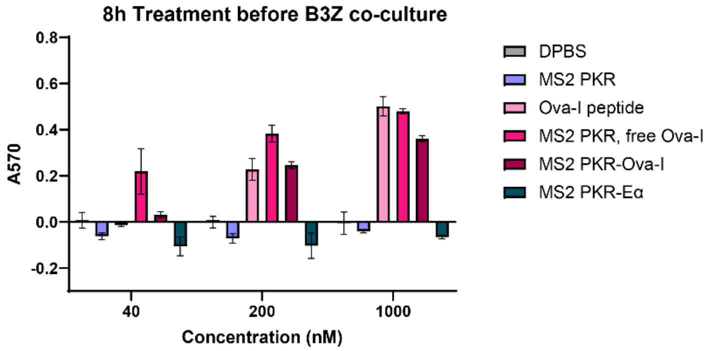


***Figure S10.*** Activation of B3Z cells co-cultured with DC2.4 cells that were treated with vaccine constructs and controls for 8 h.
